# Acute but not inherited demyelination in mouse models leads to brain tissue stiffness changes

**DOI:** 10.1101/449603

**Authors:** Dominic Eberle, Georgia Fodelianaki, Thomas Kurth, Anna Jagielska, Stephanie Möllmert, Elke Ulbricht, Katrin Wagner, Anna V. Taubenberger, Nicole Träber, Joan-Carles Escolano, Robin Franklin, Krystyn J. Van Vliet, Jochen Guck

**Affiliations:** Center for Regenerative Therapies, TU Dresden, Dresden, Germany; Institute for Clinical Chemistry and Laboratory Medicine, TU Dresden, Dresden, Germany; Massachusetts Institute of Technology, Cambridge, MA USA; Biotechnology Center, TU Dresden, Dresden, Germany; Wellcome Trust-Medical Research Council Stem Cell Institute, University of Cambridge, Cambridge, UK

**Keywords:** multiple sclerosis, tissue stiffness, AFM, demyelination, shiverer, cuprizone

## Abstract

The alteration or decrease of axonal myelination is an important hallmark of aging and disease. Demyelinated axons are impaired in their function and degenerate over time. Oligodendrocytes, the cells responsible for myelination of axons, are sensitive to mechanical properties of their environment. Growing evidence indicates that mechanical properties of demyelinating lesions are different from the healthy state and thus have the potential to affect myelinating potential of oligodendrocytes. We performed a high-resolution spatial mapping of the mechanical heterogeneity of demyelinating lesions using Atomic Force Microscope enabled indentation. Our results indicate that the stiffness of specific regions of mouse brain tissue is influenced by age and degree of myelination. Here we specifically demonstrate that acute but not inherited demyelination leads to decreased tissue stiffness, which could lower remyelination potential of oligodendrocytes. We also demonstrate that specific brain regions have unique ranges of stiffness in white and grey matter. Our *ex vivo* findings may help the design of future *in vitro* models to mimic mechanical environment of the brain in healthy and disease state. Reported here, mechanical properties of demyelinating lesions may facilitate novel approaches in treating demyelinating diseases such as multiple sclerosis.

## Introduction

The vertebrate brain consists of various types of glial and neuronal cells. Signal transduction within the brain is based on connections of neurons via their axons. Vertebrate brains are characterized by myelinated axons, allowing saltatory conduction^1^. The alteration or decrease of axonal myelination is an important hallmark in human aging and disease. Demyelinated axons are impaired in their function and degenerate over time. Multiple sclerosis is one such demyelinating disease with chronic inflammation of certain parts of the central nervous system. As of 2013, an estimated 2.3 million people were affected worldwide with currently no known cure available^2^.

It is well recognized that beside biochemical input, the mechanical characteristics of a cell’s environment can have a profound influence on its biological properties^3–5^. Neuronal and glial cells are sensitive to mechanical input during development, in disease and regenerative states^6–9^. For instance, cellular responses of spinal cord neurons such as neurite outgrowth were found to be influenced by the substrate stiffness^10^, and recently, it has been shown *in vivo* that mechanosensing of neuronal growth cones is a critical process of axonal pathfinding during neural development^11^. Other findings indicate, that astrocytes are mechanosensitive according to the compliance of their growth substrate, suggesting that *in vivo* the stiffness of surrounding tissue might have a significant effect on them^12,13^. It has been shown, that microglia are susceptible to mechanical signals^14^ and that the mechanical mismatch between nervous tissue and neuronal implants activates glial cells, resulting in gliosis and a foreign body reaction^15^.

Furthermore, oligodendrocytes (OL), the cells responsible for myelination of axons, are sensitive to mechanical properties of their environment^16^. It has been shown that sustained tensile strain can promote differentiation of oligodendrocyte progenitor cells (OPC) into OL^17,18^. Our hypothesis is that demyelinating diseases, such as multiple sclerosis, might be characterized by altered tissue stiffness at lesion sites and that this altered stiffness has an influence on biology of OPC and OL and their potential to remyelinate axons. Notably, *in vivo* studies using magnetic resonance elastography (MRE) indicate that mechanical properties of demyelinating lesions are different from the healthy state^19–21^, but the MRE image resolution within the millimeter range is too low to draw conclusions about detailed structures of lesioned tissue.

Atomic force microscopy (AFM) - based indentation measurements enable characterization of mechanical properties of neuronal tissue at high spatial resolution^22–25^. Here, we used this technique to investigate the apparent elastic modulus of young and geriatric wild-type mouse brain. Additionally, we investigated an inherited hypomyelination (shiverer^26,27^) and an acute demyelination mouse model (cuprizone^28–31^). The shiverer mouse model is characterized by an autosomal recessive mutation within the myelin basic protein (MBP) gene^32–34^. Without MBP, the axons of these mice remain hypomyelinated from birth on. Cuprizone treatment leads to a disruption of the very active metabolism of oligodendrocytes, resulting in cell death and therefore acute demyelination in affected regions^30^. We show that age as well as degree of myelination affect brain tissue stiffness. We further show that specific brain regions have unique white to grey matter stiffness ratios. Our results specifically demonstrate that acute but not inherited demyelination affects the mechanical properties of a mouse brain.

## Methods

### Ethics Statement

All animal experiments were carried out in strict accordance with European Union and German laws (Tierschutzgesetz) and were approved by the animal ethics committee of the TU Dresden and the Landesdirektion Sachsen (approval number: 2014/7).

### Animals

Young wild-type control brain slices were prepared from C57BL/6JRj animals (10 – 20 weeks old) obtained from Janvier Laboratories (France). Aged C57BL/6JOlaHsd animals were obtained from MPI-CBG Dresden with an age of 2 – 2.3 years. Shiverer mice (C3Fe.SWV-Mbp^shi^/J), a model of inherited hypomyelination, were obtained from The Jackson Laboratory (Bar Harbor, ME USA) at the age of 7 weeks. For cuprizone treatment, wildtype C57BL/6JRj age-matched male mice (12 – 14 weeks, Janvier Laboratories) were randomized into treatment (Cuprizone, 22.5 g average weight) and control (23.7 g average weight) groups. The treatment group was fed *ad libitum* a pellet chow containing 0.2% (w/w) cuprizone (C1_4_H_22_N_4_O_2_, OpenSource Diets) for 5 weeks following a setup for minimal clinical toxicity and acute demyelination^30,31^. Control mice were kept on a normal diet for the same time. After the feeding period, mice were euthanized and brains were collected.

### Mouse brain preparation

Mouse brain was freshly isolated and kept on ice in custom made HEPES buffered extracellular solution (ECS, NaCl 136 mM, KCl 3 mM, MgCl_2_ 1 mM, HEPES 10 mM, CaC_l2_ 2 mM, Glucose 11 mM, pH 7.4). After embedding in 2.5% low-gelling point agarose (Merck KGaA, Darmstadt, Germany), samples were cut at 4°C into 400 *μ*m thick sections using a Microm HM 650V Vibratome (ThermoFisher Scientific, Waltham, USA). Brain slices were then mounted on individual cell culture dishes (34 mm diameter, TPP Techno Plastic Products AG, Trasadingen, Switzerland) using Histoacryl^**®**^ tissue glue (B. Braun Melsungen AG, Melsungen, Germany). Samples were kept on ice during the preparation procedure and transferred to room temperature during measurement. FluoroMyelin^™^ live stain (ThermoFisher Scientific, Waltham, MA USA) was used to label myelinated regions in brain slices of cuprizone and cuprizone control animals by adding 10 *μ*I to slices in 4 mL ECS buffer medium 15 min before AFM measurements.

### Atomic Force Microscopy (AFM)

AFM indentation measurements were performed using the AFM setup CellHesion 200 (JPK Instruments, Berlin, Germany). A spherical polystyrene bead (diameter: 20 *μ*m, Microparticles GmbH, Berlin, Germany) was glued to an Arrow-TL1 cantilever (*k* = 13 – 37 mN/m, NanoAndMore GmbH, Wetzlar, Germany) (Fig. 1A). The calibration of the cantilever was performed before the measurements using built-in procedures of the AFM software (thermal noise method). To probe the mechanical properties of a particular region of a brain tissue section, the cantilever was positioned over the region of interest and lowered at a speed of 10μm/sec until a setpoint of 6 nN was reached. Indentation depths did not exceed 5 *μ*m. Force distance curves were collected on grids of 100 x 100 *μ*m and 7 x 7 measurement points. Three to five grids were sampled per brain section. One section was sampled per brain region of interest per mouse. The investigated brain regions contained 3 white matter areas (cerebellum white matter, corpus callosum, striatal white matter) and 4 grey matter areas (cerebellum grey matter, cortex, striatal grey matter and substantia nigra pars compacta). The resulting force-distance curves (approach part, Fig. 1B red) were analyzed using the JPK data processing software using a Hertz / Sneddon model fit for spherical indenters to determine the apparent elastic modulus using the following equations

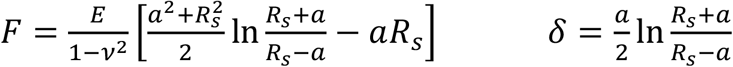

**Figure 1:**
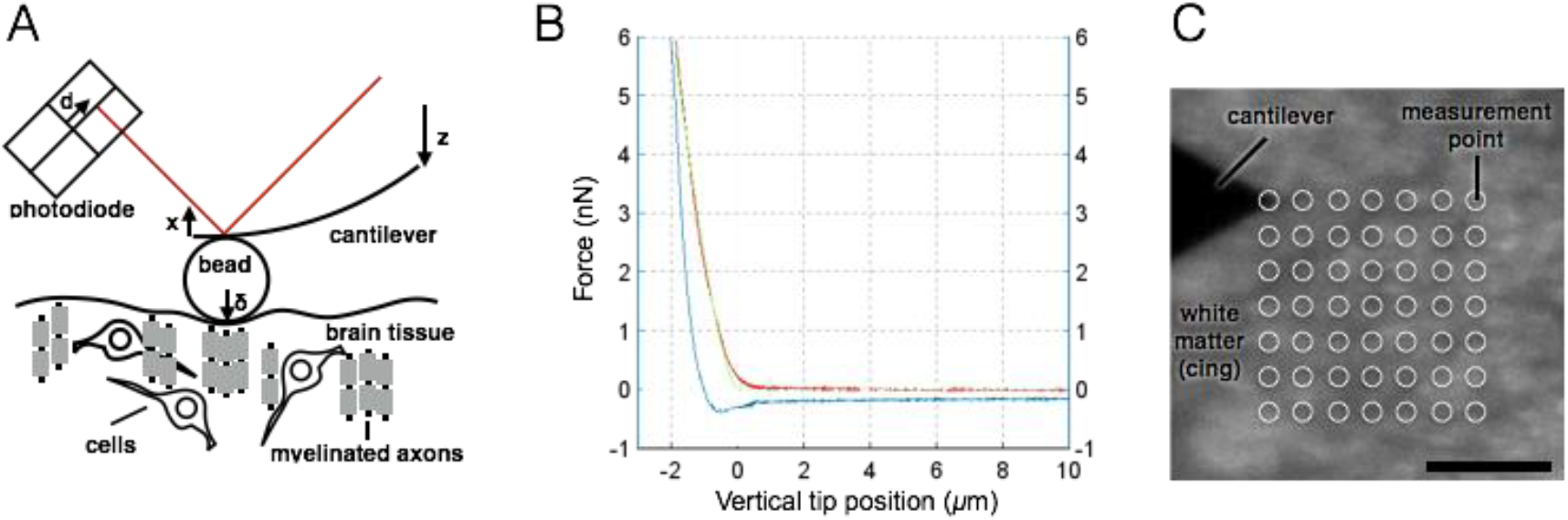
AFM measurement procedure. **A)** Schematic diagram of AFM-based indentation measurement. A spherical polystyrene bead is glued to a cantilever. A laser beam (red) directed to the backside of the cantilever is reflected and detected by a quadrant-photodiode. Upon indentation, the cantilever bends, causing a deflection of the laser beam on the photodiode. *<u>z</u>*: piezo height, x: vertical deflection of cantilever, *δ*: vertical tip position, *d*: laser deflection on photodiode. **B)** Force-distance diagram of a representative indentation. By calibrating the cantilever using the thermal noise method, the laser deflection is converted into force and is plotted on the *y*-axis. The vertical tip position (*z* – *x* in A) is plotted on the *x*-axis, resulting in a force-distance diagram. Shown is an example force curve with extend segment in red, retract segment in blue and Hertz / Sneddon model fit curve in green (JPK DP software). **C)** Example scan grid on white matter (cingulum bundle, coronal), 7 x 7 measurement points, 100 x 100 *μ*m, with approximated contact areas (white circles), scale bar: 50 *μ*m.

where *F* = force, *E* = elastic modulus, *v* = Poisson’s ratio, *δ* = indentation depth, *a* = radius of contact circle and *R_s_* = radius of sphere. We assumed the Poission’s ratio to be 0.5. ECS was always used as buffer and the temperature was kept in the range of 18 – 23°C.

### Immunofluorescent stainings

Brain slices were fixed in 4% paraformaldehyde (Sigma-Aldrich, MO, USA) overnight at 4°C. Then, the tissue was blocked for 1 hr at room temperature in PBS-Tx (1x PBS, 0.3% Triton X-100) + 0,5% Bovine Serum Albumin (BSA). Immunostaining was performed in blocking buffer at 4°C overnight using the following antibodies: rat anti-MBP (Abcam, ab7349), mouse anti-CNPase (Abcam, ab6319), sheep anti-hyaluronan (Abcam, ab53842), mouse anti-Fibronectin (DSHB, HFN7.1), rabbit anti-Iba-1 (Wako, 019-19741), rabbit anti-GFAP (DakoCytomation, Z033429-2). This was followed by washing steps in PBS-Tx and incubation with respective secondary antibodies [Cyanine-(Jackson ImmunoResearch, West Grove, PA, USA) or Alexa Fluor-conjugated (ThermoFisher Scientific, Waltham, MA USA)] in blocking solution at 4°C overnight. To stain nuclei, DAPI (4’,6-diamidino-2-phenylindole, Cell Biolabs Inc., San Diego, CA, USA) was added for 5 min. Three washing steps in PBS-Tx were done before slices were mounted on glass slides using Aqua-Poly/Mount (Polysciences Inc., Warrington, PA, USA).

Images were acquired using a light microscopic setup from Zeiss equipped with an Apotome structured illumination system (Zeiss, Oberkochen, Germany). Processing was done using Zeiss ZEN Black software and Fiji^35^, a distribution of ImageJ^36^ (v1.51g).

### Electron microscopy (EM)

For EM vibratom sections of the brain, tissue was fixed in modified Karnovsky’s fixative (2% glutaraldehyde, 2% paraformaldehyde in 50 mM HEPES) for at least overnight at 4°C. Samples were washed twice in 100 mM HEPES and twice in water, and the area of interest was cut out of the section using small pieces of a breakable razor blade. After that the samples were fixed in 2% aqueous OsO_4_ solution containing 1.5% potassium ferrocyanide and 2 mM CaCl_2_ for 30 min on ice, followed by washes in water, 1% thiocarbohydrazide in water (20 min at room temperature), again washes in water and a second osmium contrasting step in 2% OsO_4_/water (30 min, on ice). After several washes in water, the samples were *en bloc* contrasted with 1% uranyl acetate/water for 2 h on ice, washed again in water, dehydrated in a graded series of ethanol/water up to 100% ethanol, and infiltrated in epon 812 (epon/ethanol mixtures: 1:3, 1:1, 3:1 for 1.5 h each, pure epon overnight, pure epon 5hrs). Samples were embedded in flat embedding molds and cured at 65°C overnight. Ultrathin sections were prepared with a Leica UC6 ultramicrotome (Leica Microsystems, Vienna, Austria), collected on formvar-coated slot grids, and stained with lead citrate and uranyl acetate^37^. Contrasted ultrathin sections were analyzed on a FEI Morgagni D268 (FEI, Eindhoven, The Netherlands) or a Jeol JEM1400 Plus at 80 kV acceleration voltage.

### Statistical analysis

Apparent elastic modulus values were collected as grids of 100 x 100 *μ*m and 7 x 7 measurement points (Fig. 1C). Three to five grids were measured per brain region and 7 brain regions (3 white and 4 grey matter areas) were sampled per mouse. All measurement points of these grids were summarized into one brain-region-group per mouse and mean and SD values were calculated from this group. The number of data points per group is given in Table S1. Data points were processed, evaluated and graphically visualized using GraphPad Prism version 6.0h (GraphPad Software, La Jolla, CA, USA, www.graphpad.com). Unless otherwise stated, significance was tested with unpaired two-tailed t-test and given the following ratings at the graphs: ns - not significant, * - *p* ≤ 0.05, ** - *p* ≤ 0.01, *** - *p* ≤ 0.001, **** - *p* ≤ 0.0001.

## Results

### Stiffness values of different brain regions of young and old wild-type mice

No detailed examination of the elastic modulus of different mouse brain regions using AFM has been done so far. To acquire a baseline for our future experiments and to examine whether aging can affect mechanical properties of myelinated white matter (WM), we characterized the stiffness of 7 different regions of young (10 – 20 weeks) and old (100 – 105 weeks) wild-type (wt) mouse brains (Fig. 2). Three to five measurement grids were sampled per brain region and mouse, resulting in 147 – 245 measurement points per brain region per mouse. Results are given as mean ± SD and n as number of mice (see also Table S1). The investigated brain regions contained 3 WM areas (cerebellum WM, corpus callosum, striatum WM) and 4 grey matter (GM) areas (cerebellum GM, cortex, striatum GM and substantia nigra pars compacta). WM was significantly more compliant than GM in cerebellum (CB) for both, young and old mice (Fig. 2B, Table S1). No difference in stiffness was observed in young mice between cortex (CTX) and corpus callosum (cc, Fig. 2C) and WM and GM of striatum (Fig. 2D). Old mice displayed no difference between CTX and cc (Fig. 2C) but their striatal WM showed a more than 1.5-fold increase of the apparent elastic modulus, when compared with adjacent GM or WM of young mice (Fig. 2D). No significant differences were observed between the substantia nigra (SNc) GM of young and old mice (Fig. 2E).

**Figure 2:**
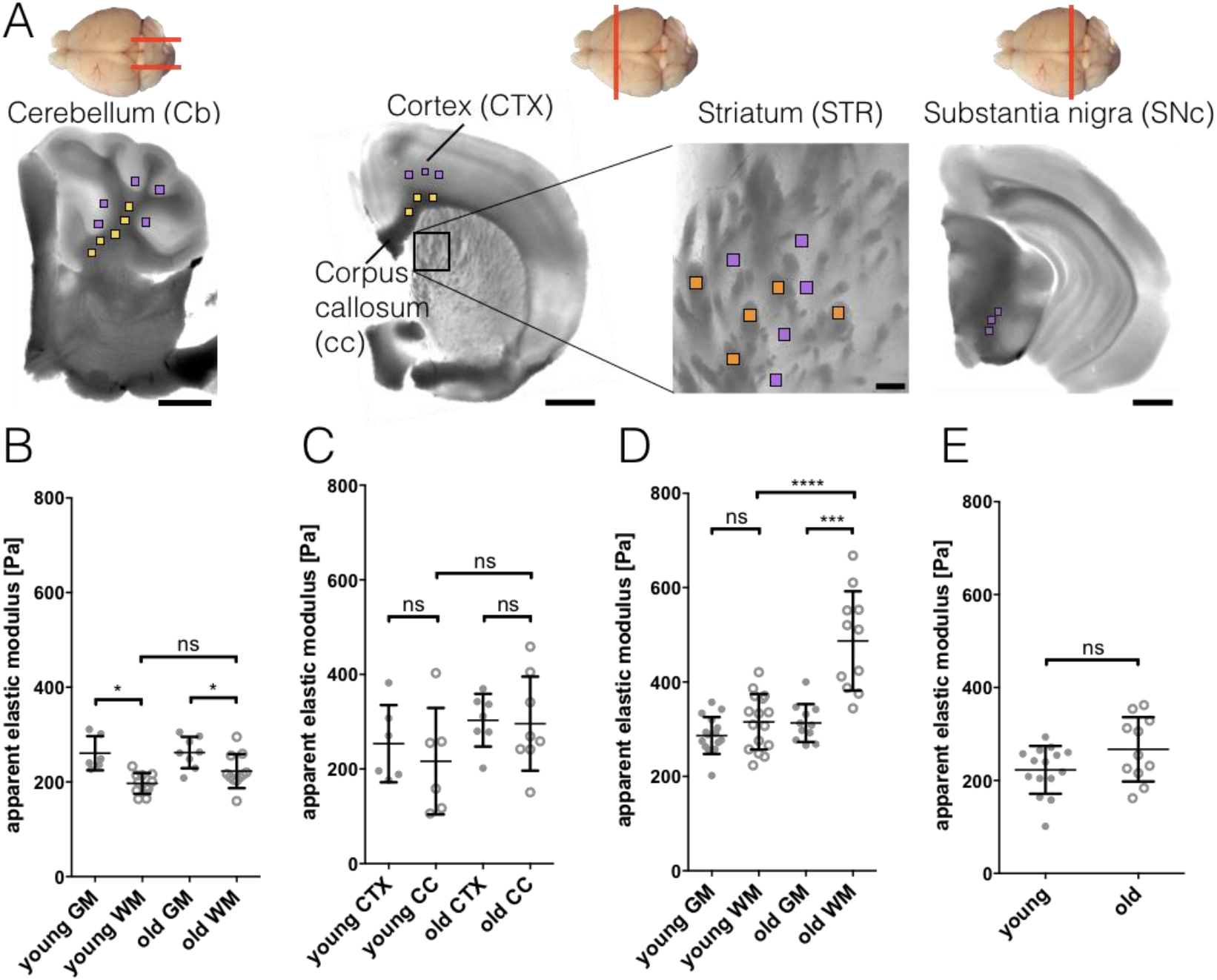
Tissue stiffness in different brain regions of young and old wild-type mice. **A)** Schematic of cutting directions of different regions of mouse brain (top view). Cerebellum: sagittal, others: coronal. Below: light microscopy images of brain regions with scan areas indicated by differently colored squares. Yellow: longitudinal WM, orange: transversal WM, purple: GM. Scan areas cerebellum (CB), cortex (CTX), corpus callosum (cc) and substantia nigra pars compacta (SNc): 100 x 100 *μ*m and 7 x 7 measurement points, scale bar: 1,000 *μ*m. Scan area striatum (STR): 50 x 50 *μ*m and 5 x 5 measurement points, scale bar: 100 *μ*m. **B-E)** Corresponding AFM-based nanoindentation results. Dots: GM, circles: WM. WM was significantly more compliant than GM in cerebellum **(B)** for both, young and old mice. No difference in stiffness was observed in young mice between cortex and corpus callosum **(C)** and WM and GM of striatum **(D)**. Old mice display no difference between cortex and corpus callosum **(C)** but their striatal WM shows a more than 1.5-fold increase of the apparent elastic modulus, when compared with adjacent GM or WM of young mice **(D)**. No differences were observed between the substantia nigra GM of young and old mice **(E)**. Data points represent means of all measurements of the respective region of individual mice. Overlay lines are mean ± SD of data points.

### Inherited hypomyelination does not affect tissue stiffness for most brain regions

The shiverer mouse is characterized by an autosomal recessive mutation within the myelin basic protein (MBP) gene. Without MBP, axons of these mice are hypomyelinated from birth on^27^. In cerebellum, corpus callosum, cortex and substantia nigra, shiverer mice showed similar values of GM and WM stiffness when compared with the same regions of young wt control mice, respectively (Fig. 3 and Table S1). Only striatal GM of shiverer mice was significantly stiffer than GM of young wt control mice (Fig. 3C), and also stiffer than corresponding striatal WM of shiverer mice.

**Figure 3:**
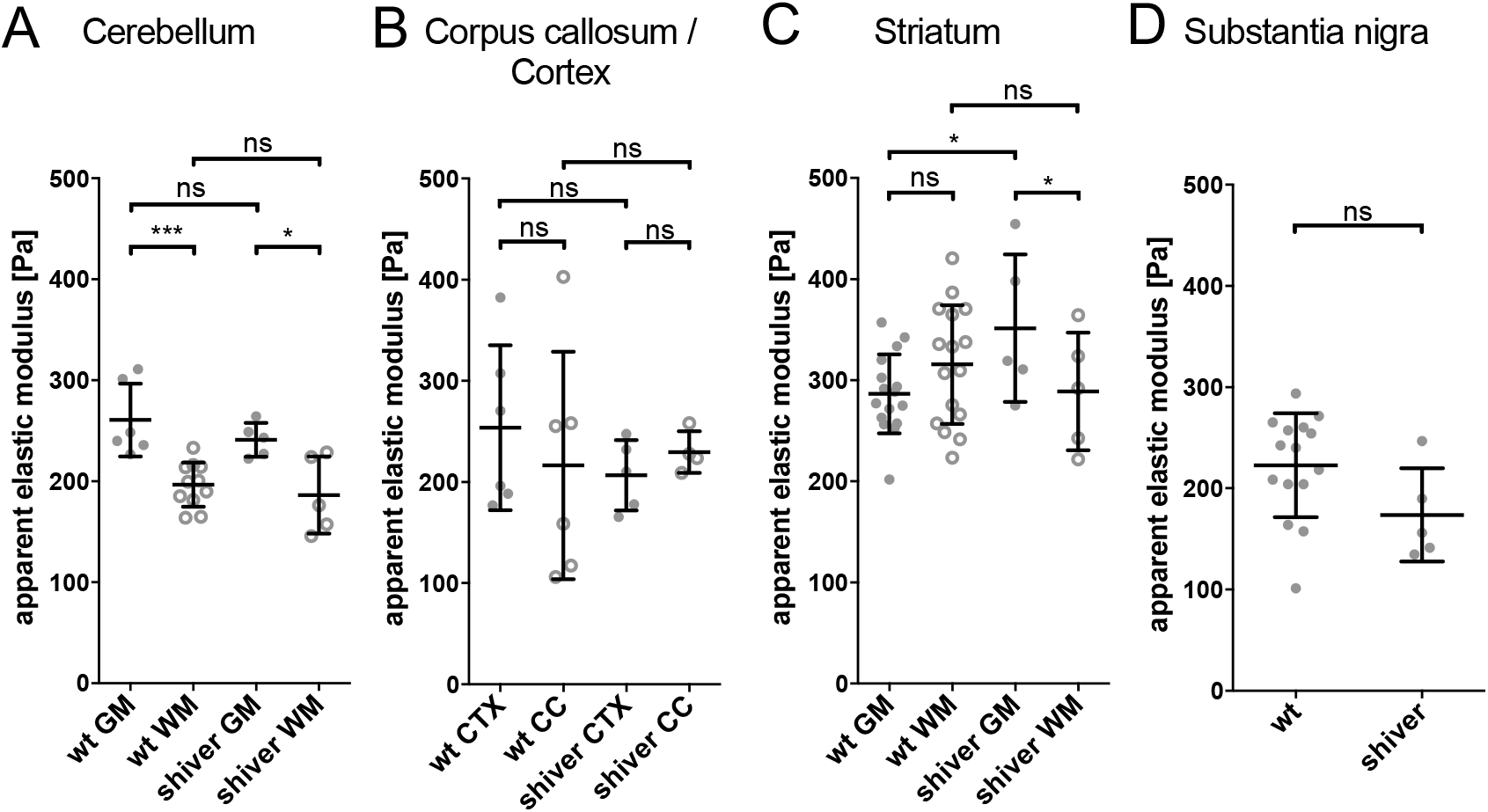
Tissue stiffness in brain regions of shiverer mice with inherited hypomyelination. In cerebellum **(A)**, corpus callosum, cortex **(B)** and substantia nigra **(D)**, shiverer mice showed similar values GM (grey dots) and WM (circles) stiffness compared to the corresponding regions in wt mice. **C)** Striatal GM of shiverer mice was significantly stiffer than GM of wild-type control mice. No significant difference could be observed between wild-type and shiverer WM. Data points represent means of all measurements of the respective region of individual mice. Overlay lines are mean ± SD of data points.

### Acute demyelination leads to decreased tissue stiffness

In contrast to shiverer mice, cuprizone-treated mice are a model of induced, acute demyelination with regionally restricted demyelinated areas^38^. Affected regions within our area of interest (corpus callosum, cc, Fig. 4A, yellow) were identified using FluoroMyelin^™^ live stain (Fig. 4B, yellow). Control measurements included the unaffected cingulum bundle region (cing, Fig. 4 A, B, orange) as well as cortex (CTX, Fig. 4A, purple). Treated animals showed markedly reduced fluorescence in certain cc regions compared to cing regions. These cc regions were chosen for AFM measurements. The acute demyelination of cc in treated mice leads to a significant 35% stiffness reduction, from 214.9 ± 19.4 Pa (control group, cc CTRL, n = 6) to 139.1 ± 16.5 Pa (cuprizone group, cc CUP, n = 8) (Fig. 4C). GM (CTX) and unaffected WM (cing) areas show no significant difference of stiffness between cuprizone treated and control groups. With 268.3 ± 36.6 Pa (CTX, CTRL, n = 7) and 271.3 ± 17.4 Pa (CTX, CUP, n = 8), GM values were approximately 55 Pa stiffer than longitudinally cut cc CTRL WM (214.9 ± 19.4 Pa, n = 6). In contrast, transversally cut, unaffected WM cing tissue (302.7 ± 72.9 Pa, n = 7 for cing CTRL and 312.5 ± 68.5 Pa, n = 8 for cing CUP) was approximately 38 Pa stiffer than GM averages.

**Figure 4:**
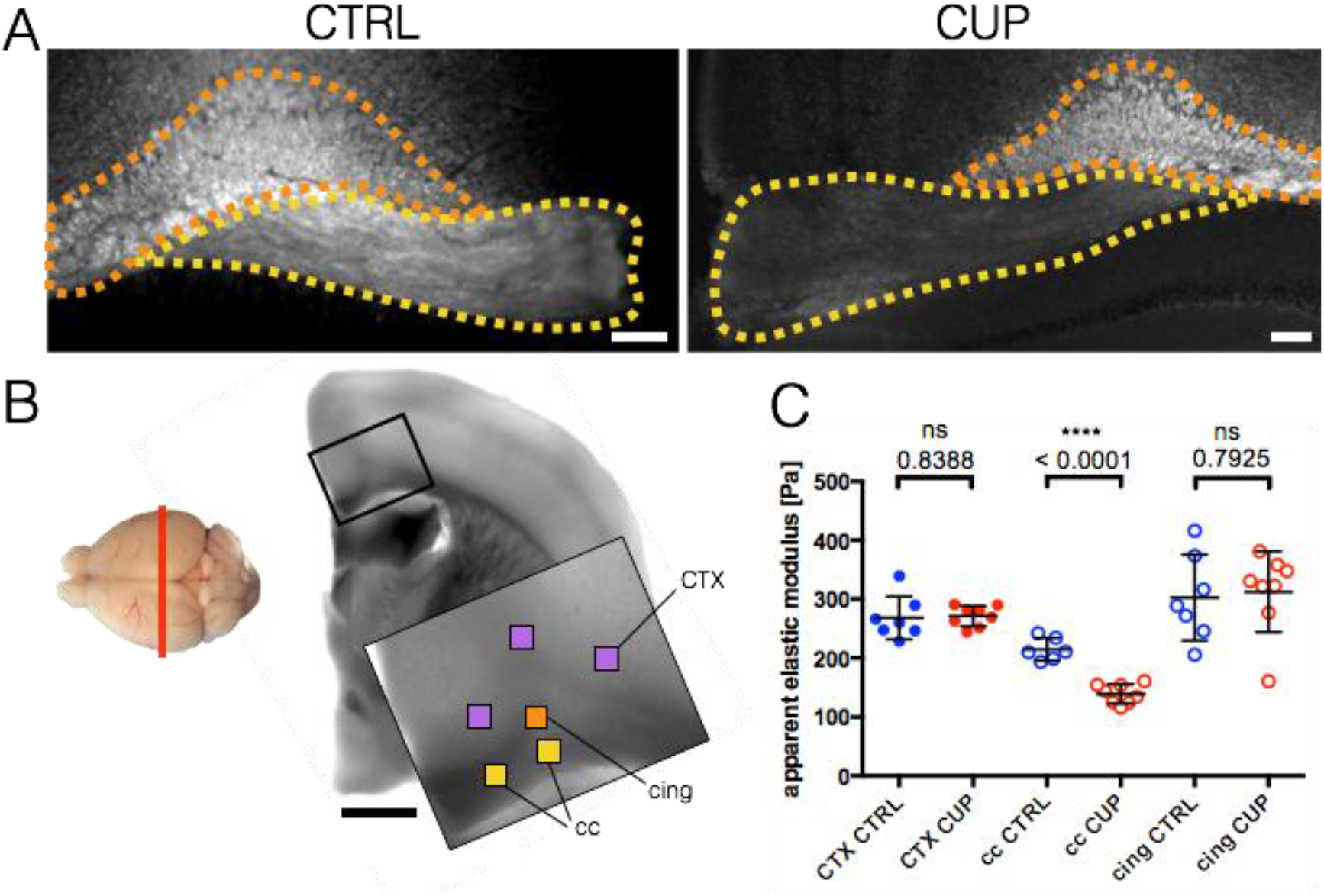
Tissue stiffness in brain regions of cuprizone treated mice with acute demyelination. **A)** Fluorescence images of control (CTRL) and cuprizone (CUP) treated corpus callosum (cc, yellow) / cingulum bundle (cing, orange) region using FluoroMyelin^™^ live stain to label myelinated regions. Treated animals showed markedly reduced fluorescence in certain cc regions compared to cing regions. These regions were chosen for AFM measurement. Scale bar: 100 *μ*m. **B)** Left side: Mouse brain top-view schematic of cutting direction (coronal). Right side: light microscopy image of brain region with scan areas indicated by differently colored squares. Yellow: longitudinal WM, orange: transversal WM, purple: GM. Scan area: 100 x 100 *μ*m and 7 x 7 measurement points, scale bar: 1,000 *μ*m. **C)** Corresponding AFM-based nanoindentation measurements. The demyelination of cc in treated mice leads to a significant drop in apparent elastic modulus of about 76 Pa on average (cc CUP). GM (CTX) and not affected WM (cing) areas show no significant difference. Dots: GM (CTX), circles: WM (cc, cing). CTRL animals labeled in blue, CUP treated animals in red. Data points represent means of all measurements of the respective region of individual mice. Overlay lines are mean ± SD of data points.

### WM stiffness differs with cutting direction

WM tissue structure is dominated by myelinated axon bundles. During sectioning of the brain, WM bundles were cut either longitudinally (Fig. S1 A, top, cc) or transversally (Fig. S1 A bottom, cing). Interestingly, when pooling all WM measurements of these two groups, transversally cut regions showed a significantly higher apparent elastic modulus (297.2 ± 86.9 Pa, n = 35) than longitudinally cut regions (210.2 ± 53.1 Pa, n = 40) (Fig. S1 B). Additionally, the standard deviation of transversally cut regions (117.1 ± 28.5 Pa, n = 23) was significantly higher than of longitudinally cut regions (84.7 Pa ± 19.3, n = 23), which might be a possible indicator of structural differences (Fig. S1 C).

## Discussion

Mechanosensitivity of cells and impact of mechanical environment on cell function have been shown *in vitro* as well as *in vivo* supporting the rising demand for systematic profiling of *in vivo* tissue mechanical properties^6–11^. Such mechanical characterization has been especially difficult for nervous tissue, because of its extremely low stiffness (elastic modulus ~1 kPa) and structural heterogeneity. Here we present for the first time a systematic high resolution (sub-micron) mechanical characterization of four distinct mouse brain regions using AFM-based indentation. This detailed mechanical profiling allowed us to compare mechanical stiffness of different regions of mouse brain and to address a question of whether mechanical changes occur in different parts of brain due to aging or disease related demyelination.

We observed relatively similar values of elastic modulus across different brain regions for GM (ranging from 200-300 Pa). Stiffness of WM showed more variation among brain regions resulting in areas where WM was more compliant than GM (CB), regions where WM was stiffer than GM (STR) and regions where there was no significant difference between grey and white matter stiffness (CTX / cc, CTX / cing). Our results agreed well with stiffness values reported earlier by Christ et al.^22^ for rat cerebellum showing WM (elastic modulus 220.5 Pa) as more compliant than GM (elastic modulus 340.5 Pa). Stiffness of WM stiffness was lower in brain slices cut longitudinally (~210 Pa) compared to those cut transversally (~ 300 Pa). This is in contrast to previous results for mouse spinal cord of Koser et al.^39^ that reported an opposite trend.

Aging-related non-mechanical changes in WM have been widely studied and are considered contributing factors to cognitive decline in older adults. During aging, myelin and nerve fibers undergo profound structural changes^40^ and remyelination efficiency declines^41–43^. Recently it has been demonstrated, that remyelination efficiency can be increased in older mice supplemented with a youthful system milieu through heterochronic parabiosis^44^. Changes in mechanical properties of CNS tissue have also been reported in rats^45^ and humans^46^ indicating an overall increase in stiffness from birth to adulthood and a decrease in stiffness with older age. However, the cause behind these stiffness changes is unknown. To further investigate the question whether age-related CNS changes, especially changes of myelin, could be linked to tissue’s mechanical properties we investigated several brain regions of young and old mice. We did not observe significant differences in stiffness between old and young mice in most of the investigated brain regions, except for the striatum. There, WM values were more than 1.5-fold increased compared to young control animals. This indicates a profound effect of aging on brain stiffness in striatum. Interestingly, this distinct region is associated with certain age-related motor function decline diseases such as Parkinson’s disease^47,48^. At the cerebellum, WM was significantly more compliant than GM for both groups. This is in line with previous results obtained for cerebellum of rats^22^.

Furthermore, we investigated CNS tissue stiffness in the inherited (shiverer) and acute (cuprizone) demyelination mouse models. While shiverer mice showed no obvious mechanical differences compared to wild-type controls, cuprizone-treated mice displayed a significant decrease in WM stiffness in affected regions characterized by significant demyelination. There could be several causes of the observed mechanical changes. First, loss of myelin in this area in cuprizone-treated mice, could lead to decreased tissue stiffness^49^. Second, oligodendrocyte death resulting in deposition of cell debris and loss of axonal connections can lead to structural changes in tissue and associated stiffness decrease (compare Fig. S2 C, C’ with D, D’). However, we and others^26,50^ observed similar structural changes in shiverer mice (Fig. S2 B, B’), which did not result in changed tissue stiffness, raising the question whether the observed structural tissue changes have a significant influence on tissue stiffness. Notably, similar brain stiffness analysis in a mouse model of tuberous sclerosis complex (TSC), which is another inherited neurological disease characterized by structurally altered brain tissue and hypomyelination, did not show stiffness difference between TSC and wild-type tissue^51^, similar to shiverer mice. Third, microglia and astrocytes proliferate and migrate to the lesion site to clean up cell debris^52^ in response to cuprizone-induced demyelination. Although increased immune cell proliferation and migration has also been reported for the shiverer mouse^50^, in direct comparison between the two models, we observed greater amounts of microglia and astrocytes in cuprizone-treated than in shiverer animals (Fig. S3, 1^st^ and 2^nd^ row), which could result in tissue stiffness changes. In general one would expect an increase of tissue stiffness when more cells per tissue area are present^39^, which is opposite to the here observed stiffness decrease in demyelinated areas of cuprizone-treated animals. However, it has been recently shown that, in contrast to other mammalian injured tissue behaviour, inflammatory responses and glial scar formation within the rodent brain, manifested by increased presence of activated astrocytes and microglia, are associated with a decrease of tissue stiffness^53,54^. It is possible that increased presence of microglia and astrocytes causes remodeling of extracellular matrix (ECM) via secreted proteins that either digest or build up ECM, which in turn may lead to changes of tissue stiffness. For example, elevated levels of ECM components laminin and collagen IV, and GFAP protein expressed in astrocytes correlate with tissue softening^54^ . Hyaluronan, a major component of ECM in the CNS, has been shown to accumulate in demyelinating lesions and inhibit rodent oligodendrocyte differentiation and remyelination^55^. Here we observed elevated levels of hyaluronan in demyelinated WM of cuprizone-treated mice (Fig. S3, last row). Because of its hydrating and viscoelastic properties^56^ hyaluronan could contribute to the observed decrease in tissue stiffness. As oligodendrocyte differentiation is enhanced on stiffer substrates^16^ it is conceivable that hyaluronan may inhibit remyelination via decrease of tissue stiffness in a lesion. Fibronectin, another component of ECM, was reported to be elevated at demyelinated lesions, which also had decreased stiffness compared to healthy CNS tissue as measured by magnetic resonance elastography (MRE)^19^. However, here we did not detect significant changes of fibronectin levels in the brain tissue of shiverer and cuprizone-treated mice (Fig. S3, 3^rd^ row).

Our results of decreased stiffness in acutely demyelinated regions agree with *in vivo* studies using magnetic resonance elastography (MRE) in rodent^19, 57^ and human brains of patients with multiple sclerosis, ^20, 21, 58^, and provide further evidence that demyelinating diseases are characterized by decreased tissue stiffness.

In summary, we demonstrate that acute but not inherited demyelination affects mechanical properties, which could contribute to a decreased remyelination potential of oligodendrocytes. It remains a subject for future studies to fully understand cellular and molecular factors contributing to tissue stiffness decrease upon acute demyelination and how these factors differ in inherited demyelination, which does not lead to similar stiffness changes.

## Acknowledgements

We acknowledge Prof. Triantafyllos Chavakis for critical reading of the manuscript, Dr. Oliver Borsch for GFAP and Iba-1 antibodies and Isabel Richter and Susanne Kretschmar for technical support. We thank the Electron Microscopy Facility and the Light Microscopy Facility (in part funded by the State of Saxony and the European Fund for Regional Development – EFRE) of the Center for Molecular and Cellular Bioengineering of the TU Dresden for imaging support. This work was supported by the National Multiple Sclerosis Society grant RG 4855A1/1 to Krystyn J. Van Vliet and by the Alexander von Humboldt Professorship grant to Jochen Guck.

## Supplemental Information

**Figure S1:**
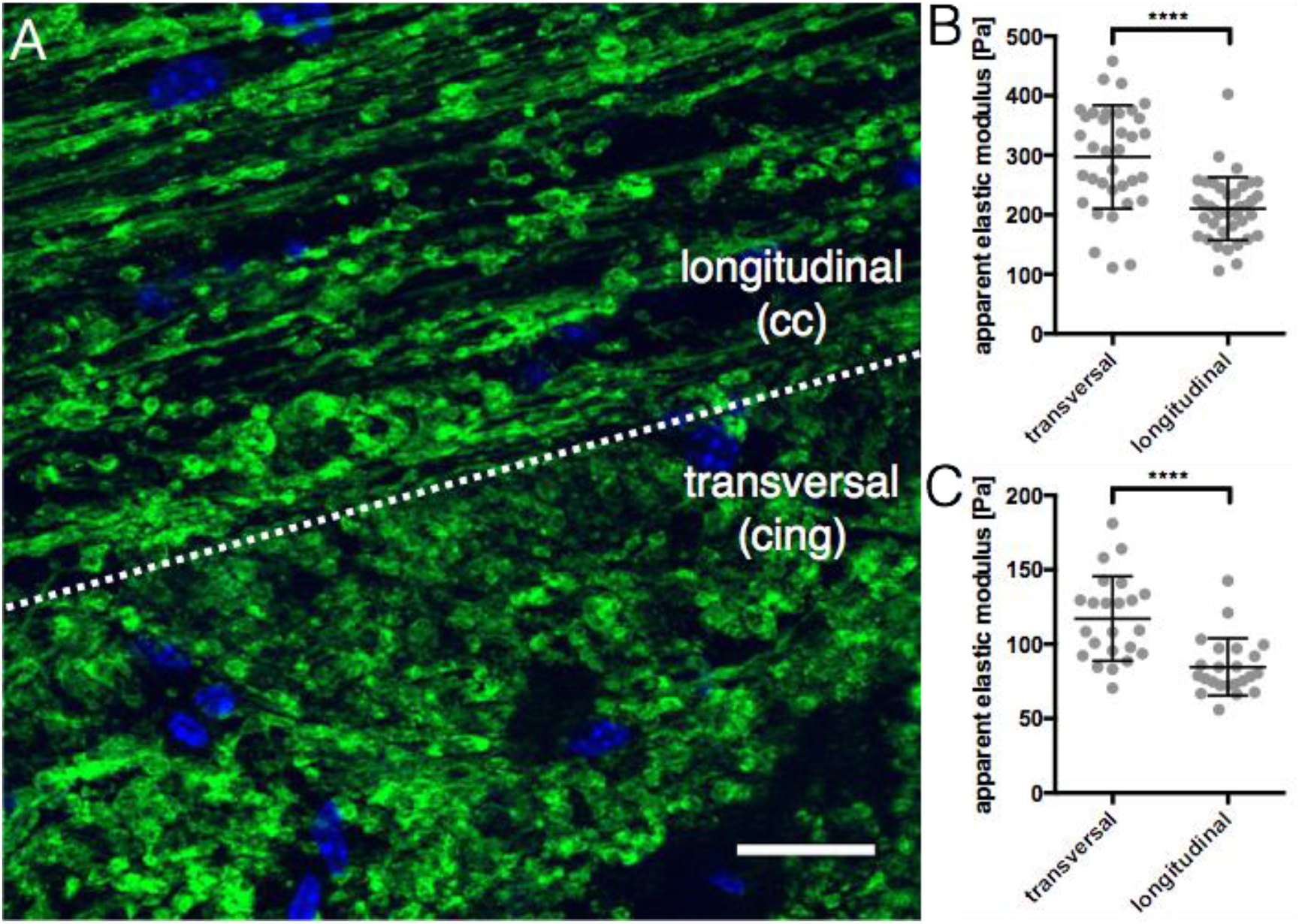
The influence of axonal direction on AFM-based nanoindentation measurements. (A) Representative image of a border region of the corpus callosum (cc) and cingulum bundle (cing) region showing longitudinally (cc region) and transversally (cing region) arranged axon bundles, stained in green using CNPase antibody. Nuclear DAPI stain is shown in blue. Scale bar: 20 *μ*m. (B) Transversally arranged WM regions (n = 35 mice) show a ~100 Pa higher apparent elastic modulus than longitudinal regions (n = 40 mice). (C) Additionally, the standard deviation of transversal regions (n = 23 mice) is significantly higher than of longitudinal regions (n = 23 mice), indicating greater structural variations. Data points represent means of all measurements of the respective region of individual mice. Overlay lines are mean ± SD of data points.

**Figure S2:**
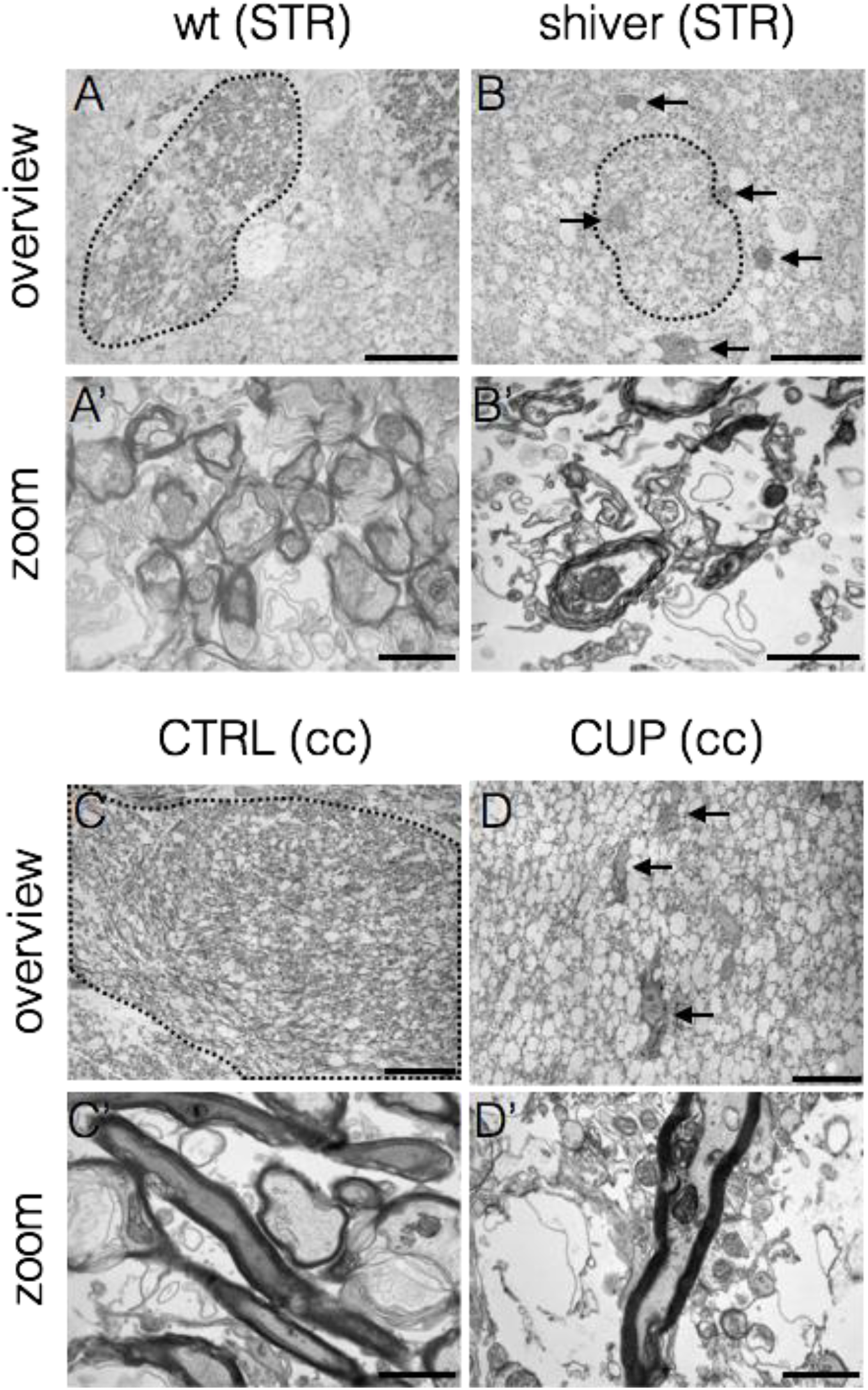
EM-Ultrastructure of WM in shiverer and cuprizone-treated mice. Wild-type (wt) mouse striatal WM (A) and cuprizone control mouse corpus callosum (C) show dense arrangements of compactly myelinated axons (A’, C’). In contrast, shiverer (B) and cuprizone-treated (D) mice show a lower amount of dark-stained myelin bundles and much looser tissue architecture (B’, D’). Additionally, dark electron-dense cells (presumably microglia, B, D, arrows) were detected in shiverer and CUP mice. These cells are characterized by a large amount of intracellular membrane vesicles, presumably phagosomes. Striatum WM (A, B) or corpus callosum (C, D) area marked with dotted black line. Scale bar: overview 20 *μ*m, zoom 1 *μ*m.

**Figure S3:**
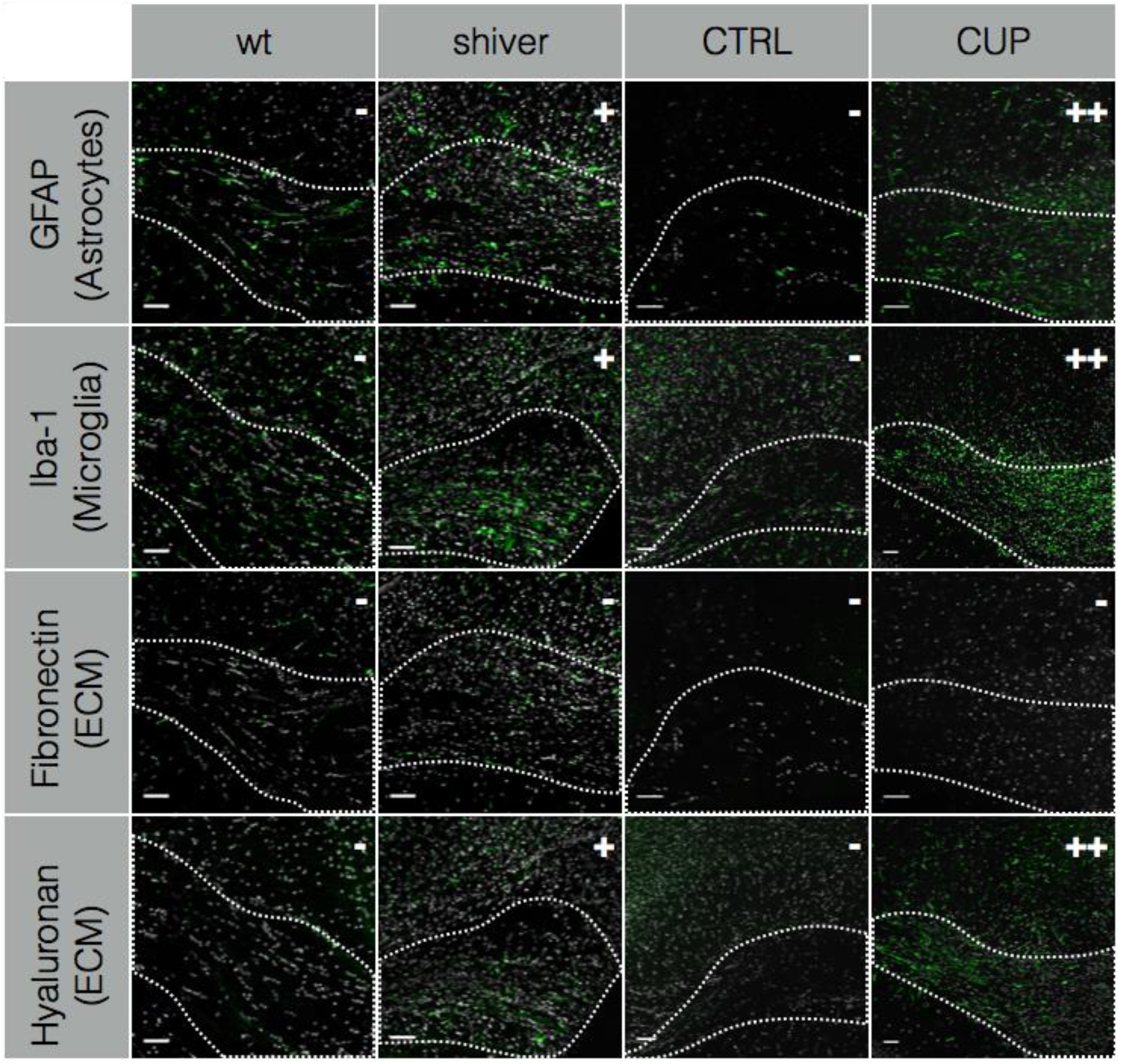
Signs of inflammation in corpus callosum of shiverer and cuprizone-treated mice. In contrast to wild-type (wt) and cuprizone control animals (CTRL), shiverer mice show more GFAP and Iba-1 positive cells and elevated hyaluronan levels. These markers increase even more in cuprizone-treated mice (CUP). Fibronectin levels remain unchanged in both, shiverer and CUP mice compared to respective controls (wt and CRTL). The respective staining (GFAP, Fibronectin, Iba-1 or hyaluronan) is shown in green, nuclear DAPI staining in white. Corpus callosum area marked with dotted white line. Staining signal intensity was categorised into (-) weak, (+) medium and (++) strong, indicated by the respective symbol in the top-right corner of each picture. Scale bar: 50 *μ*m

**Table S1:** Mean, SD and number of measurement points per experimental group. Please note that mean and SD values are calculated from means of individual mice per region and group (see also methods – statistical analysis).

| Experiment | Group | Mean [Pa] | SD [Pa] | # Mice | # Points |
| --- | --- | --- | --- | --- | --- |
| Young | CB GM | 260.6 | 36.1 | 6 | 1092 |
|  | CB WM | 196.7 | 22.0 | 11 | 2388 |
|  | CTX | 253.6 | 81.9 | 6 | 645 |
|  | cc | 216.5 | 112.5 | 6 | 702 |
|  | STR GM | 286.6 | 39.0 | 16 | 1586 |
|  | STR WM | 315.6 | 58.8 | 16 | 1643 |
|  | SNC | 222.9 | 51.3 | 15 | 1518 |
| Old | CB GM | 273.1 | 26.9 | 8 | 1492 |
|  | CB WM | 221.9 | 36.3 | 11 | 2362 |
|  | CTX | 307.6 | 54.8 | 7 | 620 |
|  | cc | 327.7 | 86.3 | 8 | 652 |
|  | STR GM | 312.2 | 39.5 | 11 | 1163 |
|  | STR WM | 504.1 | 109.0 | 11 | 1338 |
|  | SNC | 278.8 | 64.4 | 11 | 1054 |
| Shiverer | CB WM | 180.5 | 49.8 | 5 | 923 |
|  | CB GM | 239.5 | 34.1 | 5 | 876 |
|  | CTX | 206.1 | 34.7 | 5 | 378 |
|  | cc | 229.5 | 21.6 | 4 | 497 |
|  | STR WM | 286.7 | 82.67 | 5 | 493 |
|  | STR GM | 352.2 | 75.8 | 5 | 455 |
|  | SNC | 172.7 | 55.2 | 5 | 551 |
| cuprizone | CTX CTRL | 268.3 | 36.62 | 7 | 2058 |
|  | CTX CUP | 271.3 | 17.38 | 8 | 2352 |
|  | cc CTRL | 214.9 | 19.38 | 6 | 1176 |
|  | cc CUP | 139.1 | 16.47 | 8 | 1568 |
|  | cing CTRL | 302,7 | 72,86 | 7 | 594 |
|  | cing CUP | 312,5 | 68,51 | 8 | 784 |

